# Lossless Indexing with Counting de Bruijn Graphs

**DOI:** 10.1101/2021.11.09.467907

**Authors:** Mikhail Karasikov, Harun Mustafa, Gunnar Rätsch, André Kahles

## Abstract

Sequencing data is rapidly accumulating in public repositories. Making this resource accessible for interactive analysis at scale requires efficient approaches for its storage and indexing. There have recently been remarkable advances in building compressed representations of *annotated* (or *colored*) *de Bruijn graphs* for efficiently indexing k-mer sets. However, approaches for representing quantitative attributes such as gene expression or genome positions in a general manner have remained underexplored. In this work, we propose *Counting de Bruijn graphs* (Counting DBGs), a notion generalizing annotated de Bruijn graphs by supplementing each node-label relation with one or many attributes (e.g., a k-mer count or its positions). Counting DBGs index k-mer abundances from 2,652 human RNA-Seq samples in over 8-fold smaller representations compared to state-of-the-art bioinformatics tools and yet faster to construct and query. Furthermore, Counting DBGs with positional annotations losslessly represent entire reads in indexes on average 27% smaller than the input compressed with *gzip* for human Illumina RNA-Seq and 57% smaller for PacBio HiFi sequencing of viral samples. A complete searchable index of all viral PacBio SMRT reads from NCBI’s SRA (152,884 samples, 875 Gbp) comprises only 178 GB. Finally, on the full RefSeq collection, we generate a lossless and fully queryable index that is 4.4-fold smaller than the MegaBLAST index. The techniques proposed in this work naturally complement existing methods and tools employing de Bruijn graphs and significantly broaden their applicability: from indexing k-mer counts and genome positions to implementing novel sequence alignment algorithms on top of highly compressed graph-based sequence indexes.

## 1 Introduction

The sequencing of DNA and RNA has become a commodity in the portfolio of biomedical data acquisition techniques, leading to an increase both in the demand and availability of sequencing data [45]. Often, independent from the original research questions that individual data sets were created to answer, they find a second life as a valuable source for other analyses [46, 33]. Thus, methods for the efficient storage and indexing of sequence data are urgently needed. In the past years, various approaches have been proposed to address this problem. On the one side there are methods that extract relevant information, such as expression counts from vast cohorts of (RNA)-Sequencing data, and summarize it in aggregated form [13]. On the other side stand approaches that provide a full-text index of the sequencing data and allow to retrieve metadata for arbitrary sequence queries [9, 22], which is of great practical relevance for projects generating large sequence cohorts [15, 1]. As a balance between compressibility and access, methods using k-mer decompositions of the input sequences have proven very successful [36, 9, 22, 29]. In this work, we focus on approaches representing such k-mer sets as annotated sequence graphs, which we will briefly review in the following, discussing benefits and limitations of existing approaches.

### 1.1 Annotated genome graphs

To fully represent all information of a sequencing sample for interactive study, two components are necessary: i) an index representing the sequence information, allowing for the query of presence; and ii) a structure containing additional metadata, such as the biological label of a sequence or the location of a sequence within a genome context (commonly referred to as *genome coordinate*) [31, 26, 4]. Both components can be represented jointly or in separate data structures.

Conceptually, a de Bruijn graph is fully represented by its k-mer set. In practice, all k-mers must also be indexed and assigned unique numeric identifiers to allow for association with any metadata (e.g., counts or coordinates). Such indexes can be either represented explicitly as a hash table-like structure (e.g., Counting Quotient Filter [38]) or a self-index (e.g., the space-efficient BOSS representation [8]). The label information on the other hand can be encoded separately. The numeric k-mer identifiers provided by the k-mer index generate an address-space that can be used by a separate data structure holding the metadata, e.g., linking the k-mers to their presence in different input sources [32, 30, 22].

Other approaches, such as Bloom filters [12, 9, 7], do not require numeric k-mer identifiers. However, the lack of any address-space for structuring additional metadata limits their uses to only answering approximate k-mer membership queries. Thus, in this work we consider the approach encoding the metadata in a separate structure called *graph annotation* and focus on its efficient representation.

In addition, we further extend the notion of genome coordinates and introduce *k-mer coordinates*, representing the occurrence positions of a certain k-mer in the input stream. This stream may be a single genome, a list of sequencing reads, or an entire collection of arbitrary sequences. All (not only distinct) k-mers from the input are naturally ordered, and knowing this order allows reconstructing the original sequences from their corresponding paths in the graph, which we call *sequence traces*. Indeed, the first k-mer provides the first *k* characters of the first sequence, and every k-mer with the following coordinate can be used to reconstruct the next character of the sequence, while the end of the sequence can be encoded with a skipped coordinate. Hence, by representing k-mer coordinates, we encode traces of the input sequences in the graph and thereby make the index fully lossless.

### 1.2 Graph annotations

Approaches for representing relations between k-mers and input files have been extensively explored in the past decade [20, 32, 3, 23, 2, 14]. Motivated by the experiment discovery problem, which is to find a sequencing library within a large collection based on a query pattern, these methods encode binary metadata attributes (e.g., the membership of a k-mer to a certain sequence or file) in a sparse binary matrix. Depending on the number of k-mers and files, this matrix can have up to ~ 10^12^ rows (corresponding to distinct k-mers) and ~ 10^7^ columns (corresponding to different files or, in general, labels) [22]. However, it can be highly compressed thanks to its sparsity [32, 3, 23, 2, 14].

Supplementing a de Bruijn graph with this type of *binary* graph annotation provides an excellent tool for answering k-mer membership queries. However, any quantitative information of the original data is lost. In particular, queries relating to the exact occurrence position in a sequence or relating to how often the queried sequence is present in a sample can not be answered with binary annotations.

To address this problem, methods for representing non-binary graph annotations have recently started to emerge, but very few have been proposed so far. On the one end stands gPBWT, which supplements genome graphs and enables lossless encoding of haplotypes [34]. On the other end, REINDEER [30] represents approximate k-mer counts in genome graphs by averaging them within each unitig of the sample de Bruijn graphs and compressing them via run-length encoding. Unfortunately, these methods do not cover the entire spectrum of needs. In particular, REINDEER does not represent genome coordinates and does not provide a lossless representation of input sequences. In contrast, gPBWT does provide a lossless representation, but the time complexity of querying quantitative information on a pattern would be linear in the number of its occurrences, making it less well suited for indexing large read collections.

### 1.3 Sequence-to-graph alignment

Many tools have used de Bruijn graphs as indexes for alignment to collections of sequences [27, 5, 28, 43, 22, 40], applying the seed-and-extend paradigm with varying seed filtration and extension strategies. Some strategies extract sequences from the index onto which a sequence-to-sequence alignment is performed [27, 5], while others traverse the graph and compute [22] or approximate [28, 43] an alignment score. However, very few of these methods index global coordinates in reference genomes to avoid alignments to spurious paths in the graph, which would be especially helpful when aligning to complex regions in the graph with many short overlapping unitigs. To our knowledge, deBGA [27] and PuffAligner [5] (the aligner from Pufferfish [4]) are the only de Bruijn graph-based tools that index global coordinates. The more recent PuffAligner, employs a co-linear chaining approach inspired by minimap2 [26] to effectively select a good candidate location for alignment, and to limit alignment to query regions between seed hits. However, both deBGA and PuffAligner are designed for indexing long reference genomes and optimize for query performance, thus making only limited use of compression techniques and reducing their scalability. Lastly, both use k-mer hash tables to index the unitig set, restricting the minimum seed length to *k*.

### 1.4 Our contributions

In this work, we consider the problem of representing numeric attributes assigned to each k-mer–label relation in graph annotations. We call such annotations *extended graph annotations*, emphasizing that each label is supplemented with an attribute, which may include a single or multiple numeric values. In particular, we focus on indexing a) k-mer counts, representing the number of times a k-mer occurs in a certain sequencing sample, and b) k-mer coordinates, where the attributes represent all the occurrence positions of a k-mer in a sequence, a genome, or a collection thereof. Notably, the latter makes a fully lossless representation of the input sequences. Together with the underlying de Bruijn graph, such extended graph annotations make up an abstract data structure which we call a *Counting de Bruijn graph*. We demonstrate the advantage of such an index by devising a sequence-to-graph alignment algorithm called *TCG-Aligner* (Trace-Consistent Graph-based Aligner) that avoids spurious paths and correctly estimates the alignment score even when aligning sequences with repeats to loops in the graph.

## 2 Methods

In this section, we present methods and techniques ultimately employed to efficiently represent extended graph annotations in compressed data structures that cat be queried without full decompression. Assume each node-label relation (*i,j*) is supplemented with an attribute *a_i,j_*, representing a single or multiple numeric values. Naturally, such annotations can be represented as a sparse matrix, and thus, the first question to be answered is how such matrices can be represented to minimize the memory footprint, while still allowing for efficient queries without full decompression.

### 2.1 Succinct representation of sparse matrices

Here we propose a general approach for the efficient compressed representation of sparse matrices and, in particular, extended graph annotations, which supplement each binary relation kmer-label (*i,j*) with an attribute *a_i,j_* (Figure 1A, left). This attribute may be a single numeric value (e.g., the number of times k-mer *i* occurs in experiment *j*) or a set of numbers (e.g., all positions where k-mer i occurs in genome *j*). Without loss of generality, we assume a very high sparsity of the annotation matrix and decompose the initial annotation into two components schematically shown in the right part of Figure 1A: i) a binary indicator matrix representing the indexes of the entries present in the matrix, and ii) the relation attributes *a_i,j_* stored in a separate data structure, typically in form of a compressed array. The indicator matrix is then represented with a scheme supporting the rank operation on its columns or rows (depending on the layout of the attribute values), defining an ordering on the (*i,j*) pairs and enabling access to the attribute values stored in separate arrays in the consistent order.

**Fig. 1.**
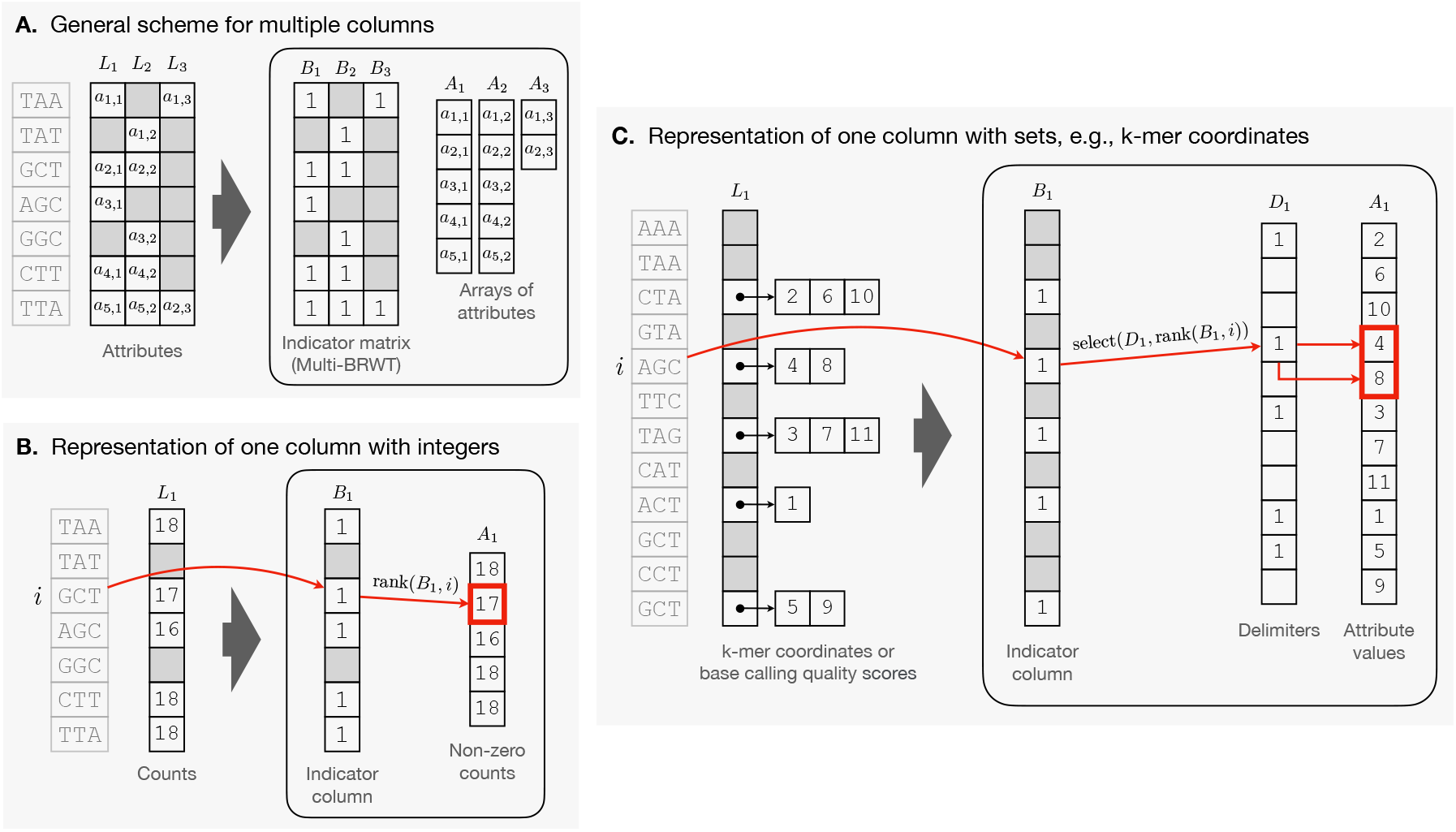
The proposed representation of sparse matrices in compressed form. **Panel A**: General scheme for sparse matrices with abstract attributes, where the non-assigned attributes are eliminated by an indicator binary matrix stored in a compressed representation (e.g., Multi-BRWT) supporting the rank operation on its columns to enable the access to the corresponding attribute for any given cell of the matrix. These attributes are stored separately, typically in a form of compressed arrays. **Panel B**: The scheme applied to a single column with integer values (e.g., k-mer counts) and the query algorithm (e.g., the count of k-mer *i* is retrieved as *A*_1_ [rank(*B*_1_, *i*)]). Empty cells in grey represent zeros. **Panel C**: The scheme applied to a single column where each cell is a set of numbers, or a tuple (e.g., representing k-mer coordinates). The “zero” attributes (empty sets) are eliminated with an indicator bitmap and the non-empty sets are encoded in an array that holds all numbers and a delimiting bitmap.

Note that the layout of the attribute arrays can be different depending on the rank operation supported by the indicator matrix. Namely, this can be the rank on its columns (shown in Figure 1), rows, or their concatenation. Thus, this scheme allows for compressing the indicator matrix using a large class of approaches for the compressed representation of binary relations, which have already been applied for representing binary graph annotations: ColumnCompressed [23], Multi-BRWT [23], BinRelWT [6], RowFlat (the scheme employed in VARI [32]), all of which support the rank operation on the non-zero entries in a certain layout.

#### Succinctness of the proposed scheme

Notably, the proposed decomposition does not change the entropy of the data, which suggests that it also does not change the theoretical minimum of the number of bits required to store the matrix by representing the two components separately.

To prove this formally, consider the problem of representing a sparse matrix of size *n* × *m* with *s* “non-zero” entries from universe 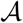 and other *nm* – *s* entries set to a fixed “zero” value that does not belong to 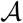. As there are 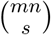 ways to pick s out of *mn* positions for non-zero values, where each can store one of 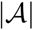 possible values, the total number of such matrices is 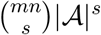. Hence, the minimum number of bits required to encode any such matrix is 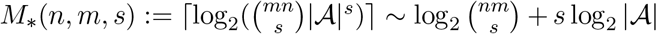. On the other hand, the indicator matrix in the proposed scheme (Figure 1A) can be reshaped into a vector encoded in the succinct Raman-Raman-Rao (RRR) representation [39] taking asymptotically log_2_ 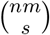 bits, which together with an optimal coding of the attributes, makes up the same space complexity ~ *M*\*(*n,m,s*). We now can make the following claim (see the proof in the Supplemental Material).

##### Theorem 1.

*If both the indicator matrix and the arrays of attributes are represented succinctly, the proposed scheme also is a succinct representation of the matrix. That is, there is no other data structure that could represent any such matrix with asymptotically fewer bits.*

#### Base representations

Now, we will show that the commonly used Compressed Sparse Column (CSC) format (e.g., used in NumPy [18]) is a special case of our proposed scheme. CSC stores the matrix-entries in three arrays: i) an array containing all non-zero values in the order they appear in the rows of the matrix, ii) an array with their column indexes, and iii) a compressed array delimiting the positions in the first two arrays corresponding to different rows of the matrix. One can see that the first array corresponds to the attribute arrays in our scheme, while the other two essentially encode the indicator matrix, making our scheme at least as efficient as the basic CSC format. Our scheme, however, allows for additional freedom in choosing specific encodings for the attribute arrays and the indicator matrix.

Encoding the columns of the indicator matrix with succinct RRR bit vector representations [39] asymptotically achieves the theoretical minimum in space. Extra compression can be achieved in practice by exploiting column-correlations using the Multi-BRWT scheme [23].

In practice, we use succinct bitmaps to represent the columns of the indicator matrix during construction. Subsequently, we convert the matrix to the Multi-BRWT representation, to reduce its final size and enhance the query speed. Typically, we compress the attribute arrays with simple bit-packing or universal coding such as Directly Addressable Codes (dac_vector) [10], supporting the direct access.

#### Representing attributes with multiple numeric values

The scheme described above effectively erases zero-entries from the matrix and stores any non-zero entries in a separate data structure (Figure 1A). We note that this approach generalizes beyond single integers as matrix entries shown in Figure 1B. In particular, it allows entries to be number sets, and hence, can be used for representing k-mer coordinates, where each k-mer may occur in multiple positions of a genome or, more generally, a collection of input sequences.

Without loss of generality, Figure 1C schematically shows how the entries of such matrices can be encoded and queried, using a single column as example. All tuples (entries) are concatenated into a single dense array storing all values together, and an additional delimiting bitmap is used to separate the different tuples. The dense array together with the delimiting bitmap represents a single array of attributes in the general scheme as shown in Figure 1A.

### 2.2 Diff-compression of extended graph annotations

Similar to the case of binary annotations, extended graph annotations often possess a certain structure that can be exploited for their compression. Indeed, attributes of nodes in the graph can often be approximated with high accuracy from the attributes of their neighbors. For example, k-mer counts, being an aggregate function of contiguous paths induced from reads, usually change incrementally, and hence, can be approximated by averaging the counts of their adjacent k-mers. Another example is k-mer coordinates, which simply shift by 1 at each node along the paths of the de Bruijn graph derived from the input sequences, which we call *traces*. Hence, one can construct an expected set of coordinates at a node if the set of coordinates at its adjacent node is known or can be reconstructed recursively.

#### Leveraging similarity of annotations of neighboring nodes

For the case of binary annotations, transformations assuming likely similarity between annotations of adjacent nodes in the graph and replacing them with relative differences have been explored in Mantis-MST [2] and RowDiff [14]. The RowDiff algorithm conceptually consists of two parts. First, for each node with at least one outgoing edge, it arbitrarily picks one of them and marks its target node as a *successor*. The subset of edges leading to the assigned successor nodes form a spanning tree of the graph. Second, it replaces the original annotations at nodes with their differences to the annotations at their assigned successor nodes. This delta-like transform is applied to all nodes in the graph except a small subset of them (called *anchors*). These anchor nodes keep the original annotation unchanged and serve to terminate every path composed of successors and break the recursion when reconstructing the original annotations (inverse transform).

Here, we devise a generalization of the RowDiff scheme to the case of extended graph annotations. We design an invertible transform, which losslessly compresses them by effectively removing the information that can be reconstructed from a neighborhood in the graph. Our generalization goes in two directions. First, in the following section, we generalize the diff-operation to act on arbitrary sets and define specific functions for the two specific cases considered in this work: k-mer counts and k-mer coordinates (genome positions). Second, we propose a more efficient algorithm for the anchor assignment (see Supplemental Section 2) and a data-driven procedure for assigning successor nodes instead of assigning them randomly as in RowDiff. The motivation of this idea is illustrated in Supplemental Figure 1 and the procedure itself is described in Supplemental Section 1. Note that this is also applicable to binary annotations and enables better compression. In addition, we consider schemes admitting the aggregation of multiple successors at forks before computing the diff (see Supplemental Section 3). This essentially replaces the diff-paths with trees and, by design, helps improve the compression at forks, where some of the traces branch out and carry their annotations away, increasing the diff.

#### Generalized diff-transform for graph annotations

Suppose the nodes in the graph are annotated with attributes from a set 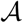. Consider a node *v* holding an attribute 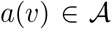 and its successor node *v_succ_* holding an attribute 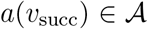. To define a diff-transform of the graph annotation, we need to specify an invertible diff-operation ⊖: 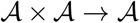 acting on pairs of attributes and replacing the original annotation *a*(*v*) with its delta relative to the annotation at the successor: *a^δ^*(*v*):= *a*(*v*) ⊖ *a*(*v_succ_*). The invertibility of this transform entails the existence of an inverse transform ⊕, such that 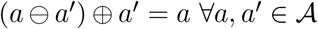, which makes it always possible to reconstruct the original annotation *a*(*v*) from the delta *a^δ^*(*v*) and the original annotation *a*(*v_succ_*), which is, in turn, either reconstructed recursively (or stored explicitly if *v_succ_* is an anchor).

For sparsifying k-mer count annotations, where the labeling at each node is encoded by a row of the integer count matrix, we use the simple vector difference as the diff operation. For the case of coordinate annotations, each attribute *a* is a set of natural numbers (occurrence positions of a k-mer in a genome or a file), that is, 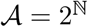. At the same time, we naturally expect the coordinates to shift by 1 when transitioning from a k-mer to its adjacent successor. Thus, the diff operation ⊖ in this case is the symmetric set difference between the coordinates at the successor node *v_succ_* and the incremented coordinates at node *v*:

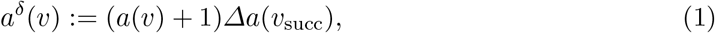

where operator *Δ* denotes the symmetric set difference: *AΔB* = (*A* ⋃ *B*) \ (*A* ⋂ *B*). Note that we chose Eq. (1) over a probably more intuitive formula *a^δ^*(*v*):= *a*(*v*)Δ(*a*(*v_succ_*) – 1) to avoid negative numbers and keep the result *a^δ^*(*v*) in the same set 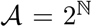 of subsets of positive coordinates.

### 2.3 Compressed extended graph annotations

In this section, we combine the techniques presented in the sections above and propose two memory-efficient representation schemes for encoding quantitative data for two important practical cases of non-binary graph annotations: k-mer counts (e.g., in read sets, representing gene expression levels) and k-mer coordinates. Coordinates may represent positions of k-mers in genomes, any collections of sequences, or in a file in general.

#### Representation of k-mer counts

Formally, the task is to represent a matrix with integer entries, where each entry corresponds to a kmer-label pair and encodes the k-mer’s count in the respective label.

We use the techniques presented above and, first, transform the initial count matrix with the generalized diff-transform (see Section 2.2). The proposed method is schematically illustrated in Figure 2. Assuming that adjacent nodes in the graph are likely to have identical or similar counts, we use the following formula to compute the diff between two rows of the integer annotation matrix: *L^δ^*(*v*):= *L*(*v*) – *L*(*v_succ_*). After this operation, the diff values *L^δ^*(*v*) are often either zeros or integer values close to zero, thus require fewer bits to represent compared to the original counts *L*(*v*). If the diff *L^δ^*(*v*) does not require fewer bits to store it than the original counts *L*(*v*) (as for nodes `GGC` and `AGC` in the example graph in Figure 2), such nodes are explicitly made anchors and no transform is applied to them (for more details, see the anchor assignment procedure described in Supplemental Section 2).

**Fig. 2.**
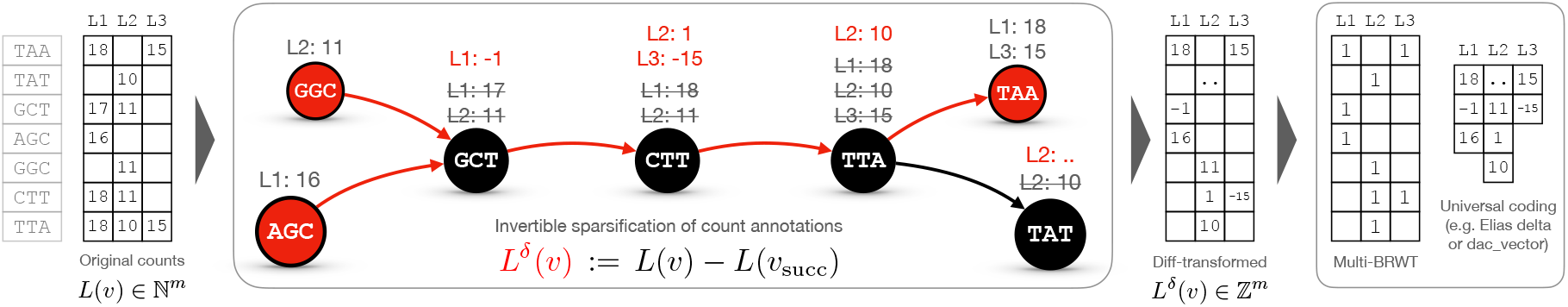
A schematic diagram illustrating encoding of k-mer counts in *m* columns with the proposed approach. Circles represent nodes of a de Bruijn graph. Edges are shown as arrows. Red nodes represent anchor nodes and red edges represent paths to row-diff successors. The transformed counts are shown in red (e.g., compare *L*_1_: –1 for k-mer `GCT` after the transform and *L*_1_: 17, *L*_2_: 11 before; for k-mer `TAT`, the transformed counts are not shown because they depend on `TAT`’s successors, not shown in the graph). Then, the diff-transformed matrix is decomposed into an indicator binary matrix stored in the compressed Multi-BRWT representation supporting the rank operation on the columns and arrays storing non-zero diffs.

Note, in work [21] developed independently in parallel to ours, a similar delta-like coding was considered for compression of k-mer counts in the framework of weighted rooted trees. This solution is close to our diff-transform for the case of a single annotation column with k-mer counts.

After performing the diff-transform, we decompose the transformed count matrix into a binary matrix of the same shape indicating the non-zero entries and a set of additional arrays containing those entries (Figure 2, right), according to the scheme shown in Figure 1A-B. The binary matrix is stored in the compressed Multi-BRWT representation and the count arrays are stored separately.

Lastly, we note that the diff-transformed values in the count arrays may be negative. We, thus, map them to non-negative integers to enable further compression with variable-length codes, using the following invertible mapping: 2(|*x*| – 1) + 1[*x* < 0], where 1[*A*] is a boolean predicate function, which evaluates as 1 if the statement *A* is true and as 0 otherwise. After this mapping, we further compress the count arrays using Directly Addressable Codes (dac_vector) [10].

#### Representation of k-mer coordinates

Finally, in this section we describe how to efficiently encode k-mer coordinates and thereby create a lossless index of the input sequences. We observe that aggregating all sequences from a single source (or label) and encoding the enumerations of each k-mer is sufficient to reconstruct the original sequences. We, thus, will consider this simplified case for brevity of the description. However, our implementation does support the explicit indexing of multiple sources with multiple annotation columns.

After all the k-mers are enumerated (Figure 3A), the underlying de Bruijn graph is annotated and the coordinates are stored in an array of lists (Figure 1C). It is easy to see that most of the adjacent pairs of k-mers are crossed by the same sequences, and hence, the successor k-mer usually has the same coordinates as its predecessor k-mer incremented by +1. Thus, it is most natural to define the diff-operation for sparsifying coordinate annotations as described in Section 2.2, Eq. (1). If this delta *L^δ^*(*v*) is an empty set, very few bits are needed to encode it. Otherwise, it would contain the information necessary to losslessly reconstruct the coordinates at the predecessor from the known (or recursively reconstructed) coordinates at the successor. This transformation step is schematically shown in Figure 3B and demonstrates how well the predictability of the coordinates at the successor nodes can be exploited to compress the coordinate annotations.

**Fig. 3.**
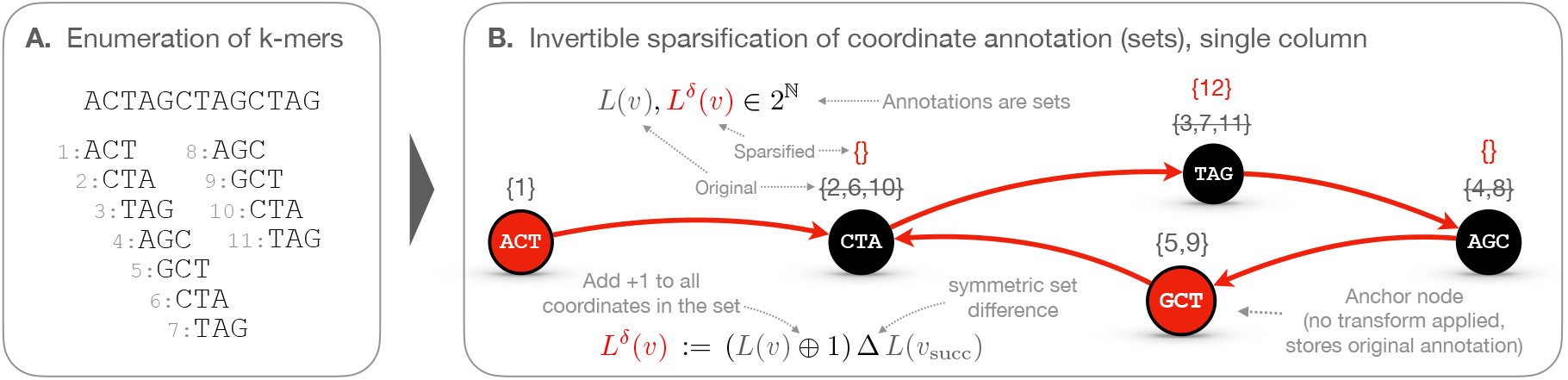
Extraction of k-mer coordinates from sequence ACTAGCTAGCTAG for k=3 (panel A) and subsequent compression with a diff-transform (panel B), where the coordinates at a node’s successor are expected to be the same but incremented by +1, as these nodes are likely to be consecutive in the input sequence(s). The symmetric set difference *AΔB*:= (*A* ⋃ *B*) \ (*A* ⋂ *B*) is used as a diff-operation. Thus, for example, *L^δ^*(`TAG`) = ({3, 7,11} ⊕ 1)Δ{4, 8} = {12}. The inverse transform is performed losslessly by *L*(*v*) = (*L*(*v_succ_*)Δ*L^δ^*(*v*)) θ 1, which follows from the following property: (*AΔB*)*ΔB* = *A* ∀*A,B*.

After the coordinate annotation is transformed, the new attributes 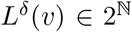 are still subsets of natural numbers, and hence, the full diff-transformed annotation matrix can be encoded with the same general scheme as shown in Figure 1C, right.

### 2.4 Sequence-to-graph alignment with k-mer coordinates

The usage of Counting de Bruijn graphs encoding k-mer counts or k-mer coordinates can greatly broaden the range of problems to which de Bruijn graphs are currently applied. In this section, we extend the sequence-to-graph alignment algorithm introduced in MetaGraph [22]. With this, we can not only ensure that all aligned paths in the graph are trace-consistent, but can also construct seed chains (detailed below) to more efficiently select good candidate positions for alignment and to reduce the number of base pairs from the query which need to be aligned.

We start by generating the initial seed set for a query sequence *q*. For low-error reads, these seeds consist of all maximal unique matches of the query which are contained in graph unitigs (called uni-MEMs [27]) with a minimum length of 19, as in [22]. For error-prone reads, we use all matches of length 19 as the seeds. For cases where the k-mer size is greater than 19, all matches are made to the suffixes of k-mers [22].

We then use the coordinates to discard all seeds which are not contained in a graph trace. Each remaining seed is associated with one or more coordinate ranges. Then, we apply a dynamic programming seed chaining algorithm similar to the approaches from Minimap2 [26] and PuffAligner [5] (see Supplemental Algorithm 2) to produce an initial partial alignment composed of a sequence of seeds (a *chain*). Let 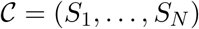 be the highest-scoring chain. For a given seed *S_i_* matching *ℓ_i_* characters with initial node *v_i_*, we denote the corresponding position in the query by *y_i_*. We complete the alignments between each pair *S_i_* and *S*_*i*+1_ by extending *S_i_* using a modification of the extension algorithm from MetaGraph [22] on the region of the query from *y_i_* to *y*_*i*+1_ + *ℓ*_*i*+1_. To complete the alignment of *q*, we extend *S_N_* forward and *S*_1_ backward until the end and beginning of *q*, respectively.

Our modification of the extension algorithm ensures that paths traverse along the corresponding graph trace of the starting coordinates *L*(*v_i_*) (in practice, only a subset originating from the top labels detected among the seeds is used for faster alignment). More precisely, we construct a trace-consistent alignment tree 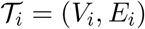 rooted at *v_i_* during graph traversal similar to the one defined in MetaGraph [22], where 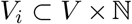 contains all the nodes along the traces originating at *v_i_*:

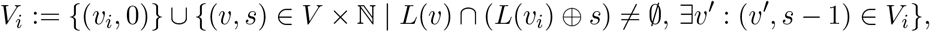

and *E_i_*:= {((*v, s*), (*v*’, *s* + 1)) ∈ *V_i_* × *V_i_* | (*v, v*’) ∈ *E*} ⊂ *V_i_* × *V_i_* contains all the edges within these paths. To avoid querying coordinates on each traversal step, we collect all reachable nodes during the forward pass of seed extension with their adjacent edges. Then, we discard non-coordinate consistent paths to obtain 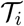 at forks in the graph (i.e., nodes with outdegree > 1). Finally, during the backtracking pass, we only extract the highest-scoring coordinate-consistent alignment (i.e., a path along 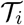) to correct for cases where the alignment terminates before the next fork node is reached. With this, we can reduce the alignment search space by more effectively filtering seeds, refining the traversal search space, and by only performing alignment on defined substrings of the query.

### 2.5 Implementation Details

The methods presented in this work were implemented within the `MetaGraph` framework https://github.com/ratschlab/metagraph with basic compression algorithms and data structures from the `sdsl-lite` library [17]. The resources and scripts used to create figures and start experiments presented in Section 3 are available at https://github.com/ratschlab/counting_dbg.

## 3 Results and Discussion

### 3.1 Indexing k-mer counts in 2,652 RNA-Seq read sets

For comparing our approach to the current state of the art, we used a set of 2,652 RNA-Seq read sets from different human tissues that was originally composed by [44] and has since been widely used for benchmarking methods indexing raw sequencing data [44, 38, 2].

We counted all 21-mers in each read set with KMC3 [24] and extracted *canonical* k-mers (defined as the lexicographical minimum of a k-mer and its reverse complement) occurring at least a certain number of times, using frequency thresholds from [38]. 66 out of the 2,652 read sets contained only reads shorter than *k* = 21 and, hence, could not be indexed. The remaining 2,586 read sets resulted in a de Bruijn graph with a total of 3.9 billion canonical k-mers and an annotation matrix of density 0.27%.

We compared the Counting de Bruijn graph to REINDEER [30], which, to the best of our knowledge, is the only published tool for indexing collections of samples with k-mer counts. We also compared against two state-of-the-art methods limited to binary annotations: Mantis-MST [2] and RowDiff [14]. The latter was used to highlight the effect of modifications made to the diff-transform presented in this work.

The results for all methods are summarized in Table 1. When compressing binary data, our approach achieved a 4.5- and 5.5-fold size improvement over Mantis-MST and REINDEER, respectively. Compared to the original RowDiff scheme, it achieved a 20% improvement in annotation compression, which resulted in an index-size reduction from 7.7 GB to 6.6 GB, thanks to our methodological improvements. The advantage is maintained when indexing k-mer counts. Applying local neighborhood-smoothing of counts along the unitigs of single-sample de Bruijn graphs, as introduced in REINDEER, our approach reduces the state-of-the-art index size of 59 GB to only 11 GB. This effect becomes even more pronounced when querying, as REINDEER needs to inflate its index for access. As we maintain our representation compressed at all times, we achieve an 8-fold reduction in memory usage during query. Even when the smoothing step is omitted, which is not possible in REINDEER, our index takes only 21 GB, while performing a lossless compression of the full k-mer spectrum of the input. Notably, our indexing workflow is also the fastest among all the compared methods in all indexing scenarios (see Supplemental Table 1) thanks to its careful implementation and parallelization, as well as leveraging the efficiently implemented KMC3 [24] and MetaGraph [22] tools.

**Table 1.** Comparison of the state-of-the-art methods for indexing raw sequencing data and the proposed approach in three scenarios of indexing 2,586 RNA-Seq read sets: i) encoding k-mer presence/absence only (binary); ii) encoding k-mer counts averaged over unitigs for each read set (smooth counts); and iii) encoding the original k-mer counts (raw counts). Query time and peak RAM were measured while querying 100 random human transcripts (≈ 90 kpb in total). If a method was not applicable to a given annotation scenario, the table shows ‘-’.

| Method | Index size |  |  | Peak RAM during query |  |  | Query time |  |  |
| --- | --- | --- | --- | --- | --- | --- | --- | --- | --- |
|  | binary | smooth counts | raw counts | binary | smooth counts | raw counts | binary | smooth counts | raw counts |
| Mantis-MST | 24.9 GB | - | - | 25.1 GB | - | - | **0.8 s** | - | - |
| RowDiff | 7.7 GB | - | - | 8 GB | - | - | 8.8 s | - | - |
| REINDEER | 30.3 GB | 59 GB | - | 58.9 GB | 91.1 GB | - | 551.2 s | 813.5 s | - |
| This work | **6.6 GB** | **11 GB** | **21 GB** | **7 GB** | **11 GB** | **21 GB** | 6.7 s | **8.2 s** | **14.4 s** |

### 3.2 Indexing read sets from the SRA

**Indexing all viral HiFi reads** In this experiment, we fetched from NCBI SRA [25] all viral sequencing samples sequenced with the PacBio Single Molecule Real-Time (SMRT) technology, including many recently sequenced SARS-CoV-2 samples. Out of all 152,957 samples (accessed on 3 Oct 2021), we could download 152,884 (99.95%) successfully. We will refer to this dataset as *Virus PacBio SMRT*. Next, we filtered this set by selecting only high-fidelity read sets to ensure a low sequencing error rate (see Supplemental Section 4 and Supplemental Figure 2). This left 152,418 read sets (99.7%), which we refer to as *Virus PacBio HiFi*. Note that here and in all other experiments, the headers of the reads (sequence names) and their quality scores were removed before indexing or compressing with the tools tested, including *gzip*.

All 152,418 read sets combined contained a total of 717 Gbp (billion base pairs) and were compressed (with headers and quality scores removed) with *gzip -9* down to 112 GB, which corresponds to 1.25 bits per base pair (bits/bp), or a compression ratio of 6.4× over the ASCII coding (8 bits/bp). Far better compression of 38× was achieved by Spring [11], a specialized method for read compression. Note, neither gzip nor Spring enable search or alignment against the input data. Then, we individually constructed lossless searchable representations of the read sets with Counting de Bruijn graphs over the {`A,C,G,T,N`} alphabet with coordinate annotations, as well as BLAST databases and sparse PufferFish indexes (see Table 2). The BLAST database required on average 2.68 bits/bp, and constructing an additional MegaBLAST index on top increased its size to an average of 125.85 bits/bp, while PufferFish required on average 21.8 bits/bp. In contrast, the compression performance of Counting de Bruijn graphs was even better than *gzip -9.* They required on average only 0.54 bits/bp (57% less than 1.25 bits/bp for *gzip -9*), while at the same time being fully searchable. For a more detailed comparison of the top four methods, see Supplemental Figure 3. We also did a similar evaluation of the methods on the full Virus PacBio SMRT dataset (see Supplemental Table 3).

**Table 2.**
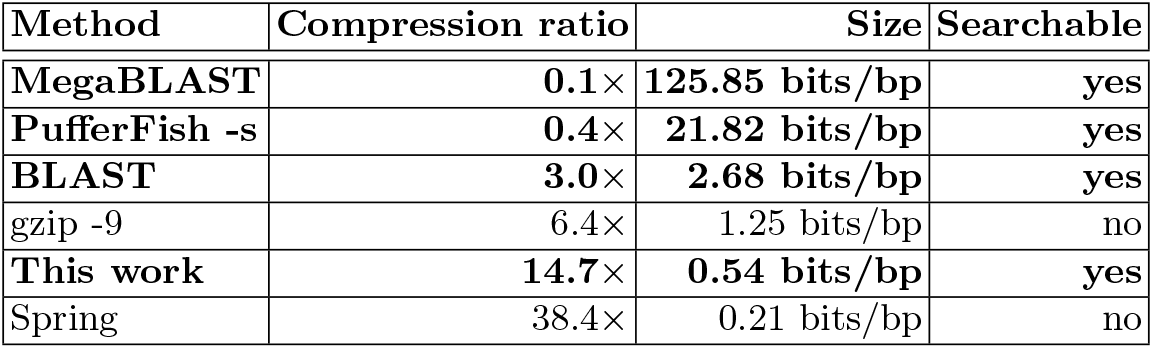
The summary of different representations constructed from Virus PacBio HiFi read sets. The methods generating searchable representations are highlighted in bold.

#### Indexing Illumina RNA-Seq reads

To evaluate compression performance on short reads, we used the RNA-Seq read sets described in Section 3.1. However, instead of indexing k-mer counts, here we constructed Counting de Bruijn graphs with coordinate annotations. We indexed all 31-mers in each read set without any other filtering. In total, 2,411 read sets (7.55 Tbp) were indexed, with the remaining samples discarded due to containing reads of variable length or only reads shorter than 31.

Again, we compared the presented approach with two alternatives commonly used for compressing read data: the general-purpose compressor gzip and the domain-specific Spring. The results are presented in Supplemental Figure 4 and discussed in Supplemental Section 5. Notably, Counting de Bruijn graphs generated on average 27% smaller representations of the input reads compared to gzip -9 (1.488 bits/bp vs. 2.030 bits/bp, see Supplemental Table 4).

#### Dependence on k-mer length

To investigate the relationship of k-mer length on index size, we compressed several representative read sets for 12 different k-mer lengths (Supplemental Figure 5). We chose one of the human RNA-Seq samples (SRR805801), representing low-error Illumina sequencing, and a *Sinorhizobium* genome sequencing sample, representing higher-error PacBio long-read sequencing. While for short, low-error reads the graph size slightly increases with *k*, the number of paths in the graph grows as well, which makes the annotation size drop steadily, due to longer unitigs length. As a result, the overall index size (combining graph and annotation) decreases with *k*. In contrast, the higher error rate in PacBio reads (SRR3747284) leads to a very large number of k-mers in the graph, and hence, its size. However, long reads with lower error (SRR13577847, PacBio HiFi) benefit from an increase of *k*. As a practical consequence, the choice of a large *k* is beneficial for compression in most scenarios.

### 3.3 Lossless index of RefSeq with k-mer coordinates

To demonstrate the use of Counting de Bruijn graphs with coordinate annotations for indexing reference genomes, we used a dataset consisting of all 32,881,422 reference sequence accessions from Release 97 of the NCBI RefSeq database [35]. Each sequence has been annotated with its associated accession ID along with all k-mer coordinates (*k* = 31). This approach forms an alternative to the commonly used MegaBLAST search tool, which requires an additional database index [31] for competitive high-throughput search. The summary of both indexes is presented in Table 3 (an extended version is available in Supplemental Table 5). Using our method, the input of 1.7 Tbp was losslessly represented in a self-index comprising 533 GB, which is 4.4× smaller than the corresponding MegaBLAST index (Table 3). We also present the performance for the subset of RefSeq containing all Fungi genomes, which we have used for demonstrating sequence alignment in the following section.

**Table 3.**
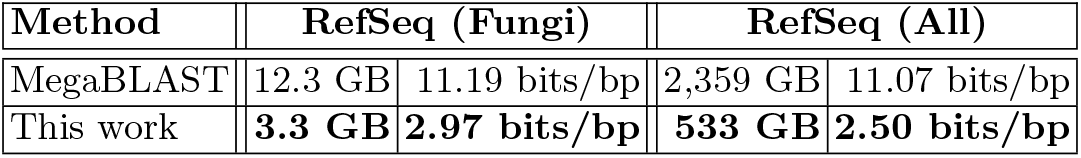
Lossless indexing of RefSeq (rel. 97) with k-mer coordinates for the complete data set (32,881,422 accessions, 1.7 Tbp) and the set of all Fungi (69,034 accessions, 8.8 Gbp).

### 3.4 Sequence-to-graph alignment with k-mer coordinates

We evaluated the accuracy of our algorithm for alignment to Counting de Bruijn graphs and compared it to the state-of-the-art aligners. We used simulated reads as queries and defined the desirable ground truth for these alignments as the corresponding segments of the reference from which the query reads were simulated. Given the human GRCh38 reference chromosome 22 [42] and the *E. coli* `NC_000913.3` [41] genomes, we use ART [19] and pbsim [37] to simulate 2000 Illumina-type and 200 PacBio-type reads of lengths 150 and 10,000, respectively. After aligning each read back to their respective reference (or graph for sequence-to-graph aligners), we compute the edit distance of the matching sequence (alignment) to the ground-truth sequence (i.e., the original reference segment from which the read was simulated) to measure alignment accuracy. In our evaluation, all nucleotides which are clipped from a query sequence by the aligner contribute as edits when measuring distance. Note, this measure is agnostic to the choice of the scoring approach used by the aligners. In addition, even a simulated read with a large number of errors can still contribute an edit distance of 0 in the evaluation if an aligner matches it to the exact ground-truth sequence.

We computed the measure described above and compared our TCG-Aligner (see Section 2.4) to other state-of-the-art methods [22, 26, 16, 5, 31, 40], run with default settings. As shown in Figure 4, the degree to which incorporation of coordinates in the alignment procedure improves accuracy and query execution time (see also Supplemental Table 2) is dependent on the complexity of the target genome. This is evident for the alignments of simulated *E. coli* reads, where the use of coordinates provides a limited improvement in accuracy and query time due to the simplicity of the genome (see the top row of Figure 4 and Supplemental Table 2). On the other hand, as can be seen in the bottom row of Figure 4, it significantly improves alignment accuracy and query time for human reads. 40.2% of human PacBio reads align to the exact ground-truth sequence when using TCG-Aligner, compared to 0.49% with the MetaGraph aligner.

**Fig. 4.**
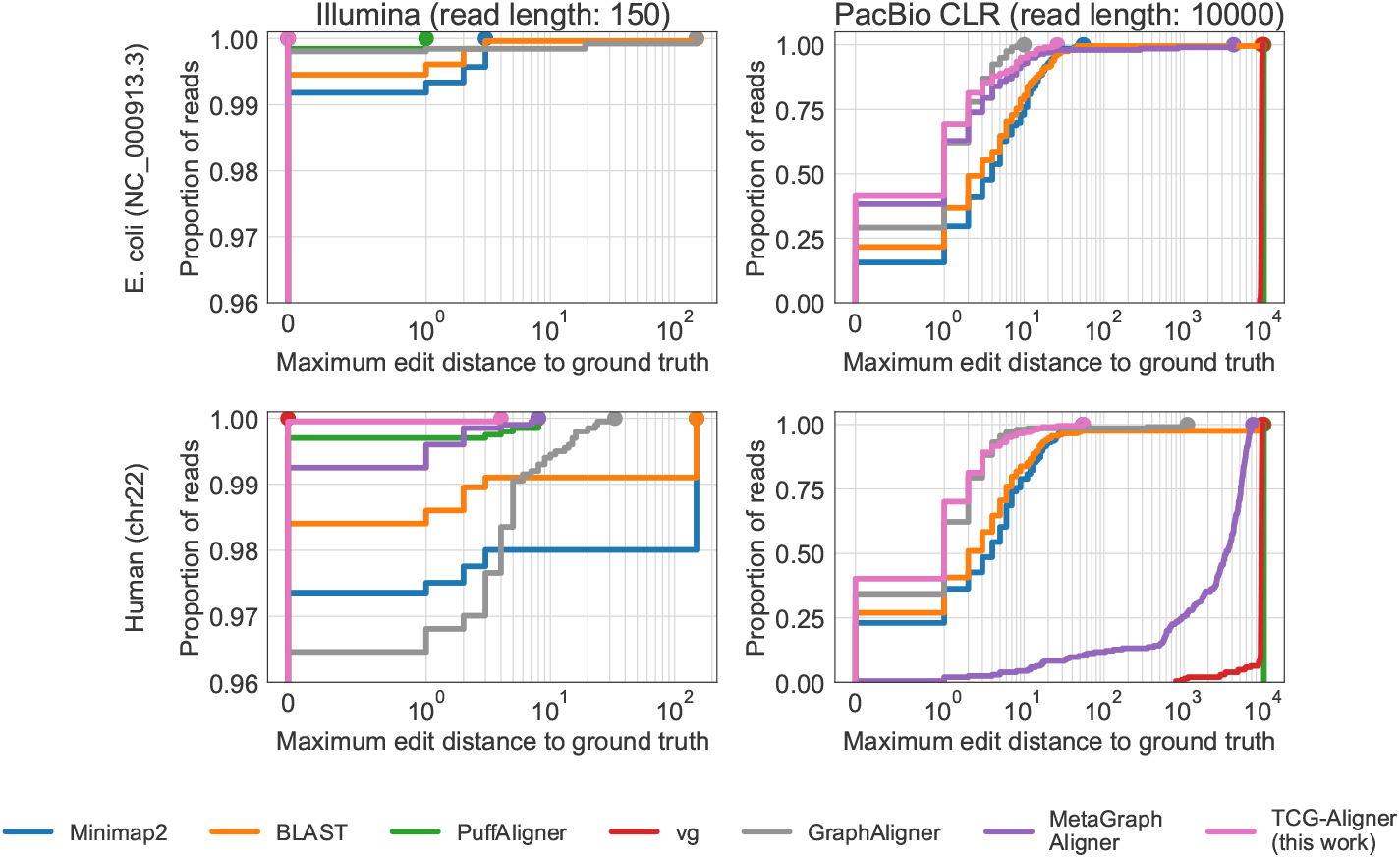
Alignment accuracy on simulated Illumina- and PacBio-type reads (*E. coli* `NC_000913.3` and human chr22). The edit distance is measured between the alignment (the returned path in the graph) and the groundtruth sequence. In the top left subplot, the curves of vg- and TCG-Aligner are superimposed.

For MetaGraph and vg, the large edit distances of the PacBio read matches relative to the ground truth are due to the aligners reporting shorter local alignments, rather than alignments of the full reads. By incorporating coordinates to more effectively filter seeds, and by restricting the alignment search space to graph traces, the TCG-Aligner provides improved accuracy and query execution time when aligning against reference genomes.

### 3.5 Searching for the Delta variant of SARS-CoV-2 in SRA

Finally, to enable fast search in the entire collection of all 152,884 viral PacBio SMRT read sets from SRA (see Section 3.2), we also indexed them in a joint Counting de Bruijn graph with coordinate annotation. Being 178 GB in size and only 19% larger than the input reads without quality scores compressed with *gzip -9* (150 GB, 875 Gbp), our index provides a fully lossless representation of them and can be used for search and alignment of arbitrary sequences. Notably, 152,272 (99.6%) of the indexed read sets originate from BioProject `PRJNA716984`, consisting of PacBio Sequel II sequencing runs from SARS-CoV-2 samples.

We queried DNA sequences flanking the nine defining mutations of the SARS-CoV-2 21A (Delta) variant spike protein against this joint index to retrieve all the occurrences of its specific mutations within the reads. In total, six sequences of length 59 and one of length 53 (to cover the deletion variants, see Supplemental Section 6 for a list of the sequences) were queried. Each sequence was matched to an average of 107,892 samples and 7.68 million positions in the joint index. Despite the enormous number of returned hits, the query took under 4 minutes on a single thread.

In this experiment, for a given sample, we classified it as containing SARS-CoV-2 21A if, for each of the defining mutations of the spike protein, there exists at least one read in the sample supporting that variant. If a sample is classified as such, we enumerate all reads containing any of the defining mutations. As shown in Figure 5, 90.5% of samples deposited after July 2021 contain each of the Delta variants of the spike protein, leading to a sharp growth in the number of reads containing these variants. We would like to note that this analysis is derived only from the dates on which these samples were uploaded to the SRA, hence, cannot be used to determine when these variants actually emerged. Although this metadata can be derived for further analysis, it is outside the scope of this work.

**Fig. 5.**
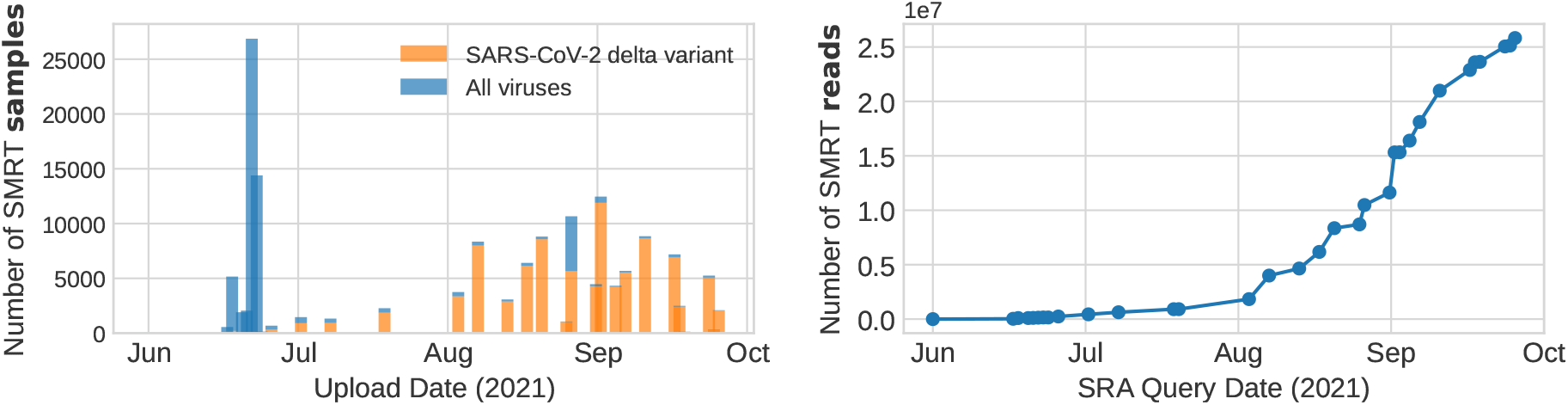
Detection of the delta variants of the SARS-CoV-2 spike protein in SMRT virus sequencing samples deposited on the SRA. **Left:** The number of samples detected to contain (orange) and not contain (blue) the SARS-CoV-2 delta variant. **Right:** The number of reads containing defining mutations contributed by samples containing the SARS-CoV-2 delta variant.

## 4 Conclusions

We have presented a novel approach for the efficient and compressed representation of quantitative annotations on de Bruijn graphs. Together with the underlying graph, these annotations make up a data structure which we call a *Counting de Bruijn graph*. It can be used to represent quantitative information and, in particular, encode traces of the input sequences in de Bruijn graphs. This not only provides a much higher flexibility of graph annotations but also allows for the truly lossless representation of any set of input sequences in them. This offers a practical solution to a long-standing problem of many methods employing de Bruijn graphs as a base structure and opens the doors to implementing novel sequence alignment algorithms on top of them. Notably, the method is agnostic to the alphabet and hence can be used not only for indexing nucleotide sequences but also protein sequences or sequences over any other alphabet.

In addition to the presented approach for the compressed representation of sparse non-binary matrices, we have generalized the RowDiff scheme [14] to non-binary graph annotations and optimized it by improving the algorithms for anchor and successor assignment. We have considered and have shown the advantages of the coding where multiple successor nodes may be assigned to each node. In future extensions, the aggregating operator *g* could act not only on the immediate successors of the node but on the whole tree of all successors spanning from it until the terminating anchor nodes. This would lead to a significantly better compression, for instance, in the case of k-mer counts linearly increasing within the paths in the graph. Moreover, an adaptive model can be trained with machine learning methods to predict annotation at a node from its successors and their annotations to further reduce the deltas stored explicitly in compressed data structures. We believe this has promising potential for this coding to benefit from the recent advances in machine learning.

Finally, we devised an algorithm for aligning sequences to Counting de Bruijn graphs with coordinate annotations, which avoids spurious paths. This algorithm correctly estimates the alignment score even when aligning sequences with repeats to loops in the graph, which would be impossible with de Bruijn graphs alone. Since sequences shared by many samples are represented by a simple path, de Bruijn graph-based approaches can greatly reduce the overhead of aligning to collections of highly similar sequences, while more traditional database search methods would align to each database entry independently. The added availability of k-mer coordinates to the MetaGraph alignment framework [22] allows for various other seeding or extension heuristics to be implemented, such as those used in MegaBLAST [31]. While the alignment method and evaluation presented here are restricted to graphs constructed from assembled reference genomes, such seed extension methods can be adapted for alignment to de Bruijn graphs constructed from raw read sets, which, we believe, is a promising direction for future work.

Playing a similar role for indexing sequences in de Bruijn graphs as gPBWT does in the realm of variation graphs and the indexing of pangenomes, we believe our approach is a significant step forward for the representation of and the search in very large collections of sequences, addressing a still increasing demand for interactive access to growing archives of biological sequences. We envision this as the first step towards enhancing the performance of BLAST-based sequence searches using graph-based approaches.

## Competing interest statement

The authors declare no competing interests.

## Acknowledgments

The authors would like to thank Mario Stanke, Matthis Ebel, Daniel Danciu, and Stefan Stark for their helpful feedback. M. K. and H. M. are funded by the Swiss National Science Foundation grant #407540_167331 “Scalable Genome Graph Data Structures for Metagenomics and Genome Annotation” as part of Swiss National Research Programme (NRP) 75 “Big Data”. M.K., H.M., and A.K. are also partially funded by ETH core funding (to G.R.).

## Author contributions

The idea and design of Counting De Bruijn Graphs and their representation techniques were conceived and developed by M. K., the TCG-Aligner was conceived and developed by H. M.. A. K. and G. R. conceptualized and supervised the research and acquired funding. All authors contributed to writing the manuscript and provided feedback on the algorithms, theory and experiments.

## Notes

### Competing Interest Statement

The authors have declared no competing interest.

### Summary of Updates

Added Supplemental Table 1 with Construction times and Peak RAM usage. Supplemental figures updated. Updated section 3.4 to clarify the setup of the alignment experiment. GraphAligner was added to the aligner comparison: Section 3.4, Figure 4, and Supplemental Table 2 updated. Also other minor corrections.

