## Supplemental Material for "Lossless Indexing with Counting de Bruijn Graphs"

### Supplemental Material:
### Lossless Indexing with Counting de Bruijn Graphs

Mikhail Karasikov[1,2,3], Harun Mustafa[1,2,3], Gunnar Rätsch[1,2,3,4,5], and André Kahles[1,2,3]

[1] Department of Computer Science, ETH Zurich, Zurich, Switzerland
[2] Biomedical Informatics Research, University Hospital Zurich, Zurich, Switzerland
[3] Swiss Institute of Bioinformatics, Lausanne, Switzerland
[4] Associate faculty in the Department of Biology at ETH Zurich, Zurich, Switzerland
[5] ETH AI Center, ETH Zurich, Zurich, Switzerland

*Proof (of **Theorem 1**).* By definition of a succinct data structure, a succinct representation of the indicator matrix takes $Z_1 + o(Z_1)$ bits, where $Z_1 = \lceil \log_2 \binom{nm}{s} \rceil$ is the minimum number of bits required to encode the matrix. Similarly, a succinct representation of the attribute vectors takes $Z_2 + o(Z_2))$ bits, where $Z_2 = \lceil s \log_2 |\mathcal{A}| \rceil$ is the theoretical minimum. Together they take $Z_1 + o(Z_1) + Z_2 + o(Z_2) = (Z_1 + Z_2)(1 + o(1)) = (\log_2 \binom{nm}{s} + s \log_2 |\mathcal{A}|)(1 + o(1))$ bits, and hence, form a succinct representation of the matrix, for which the theoretical minimum is $\lceil \log_2(\binom{mn}{s} |\mathcal{A}|^s) \rceil$.

#### 1 Optimized routing of diff-paths at forks

Even though one could make each fork node (a node with multiple outgoing edges) an anchor and store their annotations unchanged, the RowDiff algorithm went further by choosing the last outgoing edge at each node with multiple outgoing edges and extending diff-paths further without stopping at forks. Here, we notice that always taking the last outgoing edge is similar to making a random routing choice, and hence is not necessarily optimal. There are, however, cases where the routing choice would significantly change the compression performance of the transform (Supplemental Figure 1). Thus, in this work, we generalize the RowDiff scheme and allow arbitrary routings at forks (custom RowDiff successor assignments).

It is easy to see that for a locally optimal successor assignment, one needs to compute the number of bits required to encode the diff with respect to each of the possible successor nodes and take the minimum (in the example shown in Supplemental Figure 1, this optimal successor node for `CTA` would be `TAT` — case B). Formally, this procedure is described in Supplemental Algorithm 1.

This procedure essentially requires performing the diff-transform for all the nodes in the graph, which doubles the time of the whole diff-transform algorithm. Thus, we developed a heuristic algorithm for successor assignment that has a significantly lower time complexity but still drastically improves the compression ratio. The idea is based on the following observation. Forks in the graph are usually formed when a bundle of paths spanning the same nodes diverge into different directions (which usually forms bubbles or dead ends in the graph). As clearly demonstrated in Supplemental Figure 1, often the diverged paths are imbalanced in the number of logical paths constituting them: there are four paths that go from `CTA` to `TAT`, and only a single one going from `CTA` to `TAG`. Thus, selecting a successor with the largest number of annotations is usually going to be closer to optimal. We use this heuristic in practice to significantly speed up the successor assignment step of the diff-transform algorithm.

#### 2 Improved anchor assignment in RowDiff

Once the successor nodes are assigned, a set of anchor nodes has to be established to ensure a constant query time for each node. In RowDiff [1], the assignment of anchor nodes is performed in two stages. First, a set of required anchor nodes is assigned in order to break loops and satisfy the constraint on the maximum RowDiff path length (anchor assignment stage). Next, the reduction of the number of bits in each row for the RowDiff-transformed and the original matrix is computed and each node with a negative reduction is marked as an anchor (anchor optimization stage), to ensure that no row of the original matrix acquired more bits after the RowDiff transform.

Here, we notice that it is more efficient to perform these two steps in the opposite order. First, before assigning any anchor nodes, we compute for each node of the graph the expected reduction in the number of bits taken to encode the transformed annotations compared to the original annotations at the node. From this, we identify all nodes in the graph for which the diff-transform would reduce the annotation size, and those which should be anchored because transforming them would not improve the compression ratio. Next, we traverse the graph in a manner similar to the original anchor assignment algorithm from [1] and assign additional anchor nodes if needed to satisfy the remaining constraints. Already at the first step, the graph often acquires a non-trivial set of anchor nodes, some of which are also situated in diff-cycles and hence help reduce the number of additional required anchor nodes assigned at the second step.

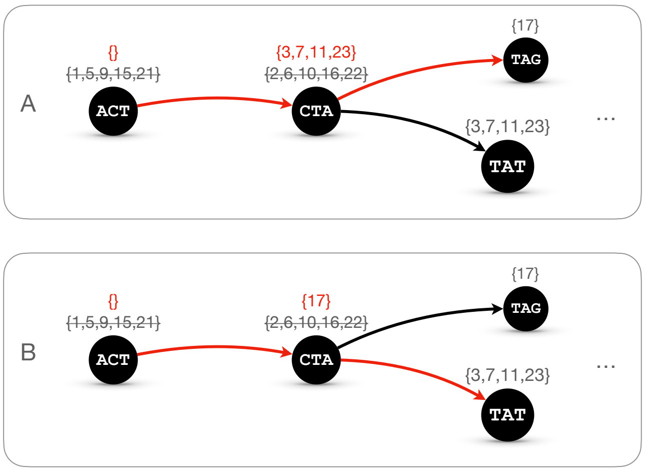


**Supplemental Figure 1.** Two cases demonstrating the advantages of the optimized routing of RowDiff paths at forks. **Panel A**: a non-optimal routing transforming the set of 5 coordinates at node `CTA` to 4 numbers; **Panel B**: the optimal routing transforming the set of 5 coordinates at node `CTA` to only 1 number.

#### 3 Assigning multiple successors at forks

Finally, we note that if we merged the annotations at both nodes `TAG` and `TAT` in Supplemental Figure 1 and computed the diff for node `CTA` against that, it would turn into an empty set, which is the best possible result. This inspires our last generalization of the successor assignment algorithm, admitting multiple successors at forks, and further improves the compression ratio (e.g., see results in Supplemental Figure 4). However, the query time also grows with the size of the successor tree, and can be exponentially large with its depth. Thus, we iteratively try to assign more successors at each fork, until the total size of the corresponding successor tree remains under a pre-defined threshold.

If node $v$ has multiple successor nodes assigned to it, their annotations $a(v^1_{\text{succ}}), \dots, a(v^t_{\text{succ}})$ are aggregated with an operator $g : \cup_{t=1}^{\infty} \mathcal{A}^t \to \mathcal{A}$ and used for computing the delta:

$$a^\delta(v) := a(v) \ominus g(a(v^1_{\text{succ}}), \dots, a(v^t_{\text{succ}})). \tag{1}$$

Note that there are no constraints on the operator $g$. Moreover, in general, this could act not only on the immediate successors of $v$, but on the whole tree rooted at $v$ and containing all successors until anchor nodes.

When sparsifying k-mer count annotations, the aggregation is performed via the simple summation of the annotation rows corresponding to the successor nodes, where the labeling at each node is encoded by a row of the integer count matrix.

For the case of coordinate annotations, each attribute $a$ is a set of natural numbers (occurrence positions of a k-mer in a genome or a file), and thus, the aggregating operator $g$ is the simple set union $g(a(v^1_{\text{succ}}), \dots, a(v^t_{\text{succ}})) = \cup_{i=1}^{t} a(v^i_{\text{succ}})$.

#### 4 Selection of PacBio HiFi read sets

The metadata indicating whether a PacBio read set consists in HiFi reads is not stored at NCBI systematically, so we selected for these read sets manually based on the sequencing quality scores. More precisely, for PacBio HiFi sequencing, the quality scores are usually dominated by symbol '$\sim$' encoding the highest possible quality score. Based on this, we applied a threshold of 0.7 for the frequency of '$\sim$' in the quality scores in the first 100 reads of a read set (see the distribution of these frequencies in Supplemental Figure 2), which resulted in 152,418 read sets, corresponding to 99.7% samples from the initial Virus PacBio SMRT dataset.

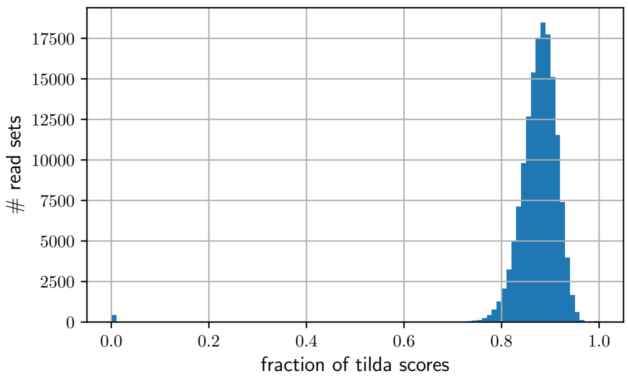


**Supplemental Figure 2.** Distribution of the frequency of '∼' in the quality scores of the first 100 reads in 152,884 Virus SMRT read sets.

#### 5 Indexing reads in Illumina RNA-Seq read sets

We indexed each of the 2,411 RNA-Seq read sets in Counting de Bruijn graphs with assigning multiple successor nodes at forks (see Supplemental Section 3). On average, Counting de Bruijn graphs are 27% smaller than the input reads compressed with *gzip -9* (1.488 bits/bp vs 2.030 bits/bp, see Supplemental Table 4). Taking a closer look at the samples showing the worst performance, we found that they typically represent functional genomics experiments, with technical sequences present in the reads (e.g., adapters in SRR808403), which can not be assembled into larger underlying sequences as expected for genome or transcriptome sequencing. Removing the adapter sequences from these reads would likely drastically increase their compressibility with Counting de Bruijn graphs.

When compared to Coutning de Bruijn graphs with a single successor node assigned per fork, our generalization with assigning multiple successors reduced the average index size by 9% (from 1.636 bits/bp to 1.488 bits/bp).

Note that similarly to all other experiments, we dropped quality scores and read headers from all inputs. In addition, here we assumed that all reads in each sample are of the same length, which helped further reduce the index size.

#### 6 SARS-CoV-2 delta variant query sequences

```
>OK091006.1:21536-25357:27-85
ACTAGTCTCTAGTCAGTGTGTTAATCTTAGAACCAGAACTCAATTACCCCCTGCATACA
>OK091006.1:21536-25357:444-502
CAACAAAAGTTGGATGGAAAGTGGAGTTTATTCTAGTGCGAATAATTGCACTT
>OK091006.1:21536-25357:1326-1384
TTCTAAGGTTGGTGGTAATTATAATTACCGGTATAGATTGTTTAGGAAGTCTAATCTCA
>OK091006.1:21536-25357:1404-1462
TTCAACTGAAATCTATCAGGCCGGTAGCAAACCTTGTAATGGTGTTGAAGGTTTTAATT
>OK091006.1:21536-25357:1812-1870
TTCTAACCAGGTTGCTGTTCTTTATCAGGGTGTTAACTGCACAGAAGTCCCTGTTGCTA
>OK091006.1:21536-25357:2013-2071
CGCTAGTTATCAGACTCAGACTAATTCTCGTCGGCGGGCACGTAGTGTAGCTAGTCAAT
>OK091006.1:21536-25357:2820-2878
CACAGCAAGTGCACTTGGAAAACTTCAAAATGTGGTCAACCAAAATGCACAAGCTTTAA
```

### 7 Supplemental Tables

**Supplemental Table 1.** Time and maximum RAM usage for indexing 2,586 RNA-Seq read sets with the state-of-the-art methods and the proposed approach in three scenarios: i) encoding k-mer presence/absence only (binary); ii) encoding k-mer counts averaged over unitigs for each read set (smooth counts); and iii) encoding the original k-mer counts (raw counts). If a method was not applicable to a given annotation scenario, the table shows '-'. The construction time represents how long it takes to construct the index on a machine with 36 physical cores Intel® Xeon® Gold 6140. Each indexing workflow was broken down into its constituting steps, for which the time and max. RAM were measured. Steps marked with '*' could only be executed with a single thread (at the time of writing, Mantis did not have an option for parallel construction, while REINDEER did but terminated with critical errors when using more than one thread). Squeakr did support using multiple threads, but turned out to be faster with 1 thread. All other construction steps for all methods were executed using 2 threads per physical core. When running Squeakr with 2 threads (see first two rows in gray in the table), we first estimated the sample's k-mer spectrum using ntCard (as recommended by Squeakr), then used the `lognumslots.sh` script packaged with Squeakr to estimate the number of slots in the counter.
Steps marked with '†' were performed by processing the 2,586 read sets in separate tasks running in parallel, using 1 core and 2 threads per task (except Squeakr, which used 1 thread per task). Construction time for these steps was computed as the sum of all wall times for all tasks divided by 36 (the number of cores and tasks running in parallel). Max. RAM was computed as the sum of RAM usages for the 36 top RAM-using tasks, which represents the maximum possible RAM usage of random task scheduling.

| | **Construction time** | | | **Max. RAM** | | |
|---|---|---|---|---|---|---|
| **Method** | **Binary** | **Smooth counts** | **Raw counts** | **Binary** | **Smooth counts** | **Raw counts** |
| k-mer spectrum estimation with ntCard† | 2.0 h | - | - | 19 GB | - | - |
| k-mer counting with Squeakr† | 35.4 h | - | - | 333 GB | - | - |
| k-mer counting with Squeakr (single thread)†* | 19.3 h | - | - | 483 GB | - | - |
| Graph and annotation construction with Mantis* | 9.6 h | - | - | 42.7 GB | - | - |
| Annotation transform to MST | 0.4 h | - | - | 26.6 GB | - | - |
| **Mantis-MST (all steps)** | **29.3 h** | - | - | **483 GB** | - | - |
| k-mer counting with KMC† | 3.2 h | - | - | 213 GB | - | - |
| Assembling contigs from k-mers with MetaGraph† | 0.7 h | - | - | 51 GB | - | - |
| Graph construction and annotation with MetaGraph | 1.7 h | - | - | 52 GB | - | - |
| Annotation transform to RowDiff-MultiBRWT | 1.3 h | - | - | 58 GB | - | - |
| **RowDiff (all steps)** | **6.9 h** | - | - | **213 GB** | - | - |
| Assembling unitigs with BCALM† | 41.5 h | 41.5 h | - | 248 GB | 248 GB | - |
| REINDEER (indexing)* | 8.7 h | 10.6 h | - | 68.8 GB | 59.4 GB | - |
| **REINDEER (all steps)** | **50.2 h** | **52.1 h** | - | **248 GB** | **248 GB** | - |
| k-mer counting with KMC† | 3.2 h | 3.2 h | 3.2 h | 213 GB | 213 GB | 213 GB |
| Assembling contigs from k-mers with MetaGraph† | 0.7 h | 0.9 h | 0.8 h | 51 GB | 70 GB | 70 GB |
| Graph construction and annotation with MetaGraph | 1.6 h | 2.2 h | 2.3 h | 52 GB | 52 GB | 52 GB |
| Annotation transformation (this work) | 1.2 h | 1.2 h | 1.4 h | 59 GB | 88 GB | 88 GB |
| **This work (all steps)** | **6.7 h** | **7.5 h** | **7.7 h** | **213 GB** | **213 GB** | **213 GB** |

**Supplemental Table 2.** Percentage of simulated reads mapping exactly to their respective ground-truth sequences (top) and total query time in seconds (bottom). For each reference, 2000 Illumina reads were simulated using ART and 200 PacBio CLR reads were simulated using pbsim, respectively. PuffAligner was unable to align the PacBio-type reads.

| **Reference** | **Minimap2** | **BLAST** | **PuffAligner** | **vg** | **GraphAligner** | **MetaGraph-Aligner** | **This work** |
|---|---|---|---|---|---|---|---|
| **Illumina**: `E. coli` | 99.1% | 99.45% | 99.84% | **100.00%** | 99.8% | **100.00%** | **100.00%** |
| **Illumina**: `chr22` | 97.36% | 98.4% | 99.70% | **100.00%** | 96.46% | 99.25% | 99.95% |
| **PacBio**: `E. coli` | 15.58% | 21.61% | N/A | 0.00% | 29.15% | 38.69% | **41.71%** |
| **PacBio**: `chr22` | 23.04% | 26.96% | N/A | 0.00% | 34.31% | 0.49% | **40.20%** |
| **Illumina**: `E. coli` | 0.13 s | 0.39 s | **0.03 s** | 1.16 s | 3.33 s | 1.13 s | 1.16 s |
| **Illumina**: `chr22` | **0.47 s** | 26.69 s | 0.96 s | 5.67 s | 44.27 s | 6.67 s | 3.89 s |
| **PacBio**: `E. coli` | **1.11 s** | 1.51 s | 4.31 s | 443.23 s | 7.60 s | 40.20 s | 35.09 s |
| **PacBio**: `chr22` | **2.43 s** | 34.80 s | 4.91 s | 1059.05 s | 50.68 s | 828.91 s | 83.48 s |

**Supplemental Table 3.** The summary of different representations constructed from all 152,884 viral PacBio SMRT read sets from NCBI Sequence Read Archive. The methods generating searchable representations are highlighted in bold.

| Method | Compression ratio | Size | Searchable |
|---|---|---|---|
| **MegaBLAST** | **0.1×** | **105.4 bits/bp** | **yes** |
| **PufferFish -s** | **0.3×** | **25.1 bits/bp** | **yes** |
| **BLAST** | **3.1×** | **2.6 bits/bp** | **yes** |
| gzip -9 | 5.8× | 1.4 bits/bp | no |
| **This work** | **6.2×** | **1.3 bits/bp** | **yes** |
| Spring | 18.0× | 0.44 bits/bp | no |

**Supplemental Table 4.** The summary of different representations constructed from 2,411 RNA-Seq read sets. The methods generating searchable representations are highlighted in bold.

| Method | Compression ratio | Size | Searchable |
|---|---|---|---|
| gzip -9 | 3.9× | 2.030 bits/bp | no |
| **This work (single succ.)** | **4.9×** | **1.636 bits/bp** | **yes** |
| **This work (mult. succ.)** | **5.4×** | **1.488 bits/bp** | **yes** |
| Spring | 12.2× | 0.658 bits/bp | no |

**Supplemental Table 5.** Lossless indexing of RefSeq (release 97) with k-mer coordinates.

| | | | | BLAST | | This work | | |
|---|---|---|---|---|---|---|---|---|
| **Data** | **# accessions** | **# base pairs** | **gzip -9** | DB only | **MegaBLAST** | Graph | Annotation | **Total** |
| RefSeq (Fungi only) | 69,034 | 8.8 Gpb | 2.6 GB | 2.2 GB | **12.3 GB** | 2.2 GB | 1.1 GB | **3.3 GB** |
| RefSeq (All) | 32,881,422 | 1.7 Tbp | 483 GB | 437 GB | **2,359 GB** | 176 GB | 357 GB | **533 GB** |

#### 8 Supplemental Algorithms

**Supplemental Algorithm 1:** Optimal successor assignment

**Input:** graph $G$, diff-operation $\ominus$, annotations $L(v)$, estimator of the number of bits required to store annotations $Size(L(v))$

**Output:** succ(u) — selected successor nodes

**foreach** *node* $u \in Nodes(G)$ **do**
  succ(u) $\leftarrow \arg\min_{v \in \mathrm{Out}_G(u)} Size(L(u) \ominus L(v))$
**return** succ

**Supplemental Algorithm 2:** Seed chaining algorithm

**Input:** seed sequence $\mathcal{S} = \{S_1, \ldots, S_N\}$, where $S_i \in \mathcal{S}$ is an exact match storing the match of the query region $[y_i, y_i + \ell_i]$ to the single reference region $[x_i, x_i + \ell_i]$. $\mathcal{S}$ is sorted by non-increasing $(x_i, y_i)$. $b$ is a bandwidth (for us, $b = 200$). $\oplus$ is the sequence append operator. $L$ is the seed length (for us, $L = 19$)

**Output:** $(\mathcal{C}, s)$ — a chain (a subsequence of $\mathcal{S}$) with score $s$

`// Initialize dynamic programming and backtracking arrays`
**foreach** $i \in \{1, \ldots N\}$ **do**
  $C[i] = \ell_i$ `// Initialize chain score`
  $B[i] = 0$
`// Forward pass`
**foreach** *seed* $S_i \in \{S_1, \ldots, S_{N-1}\}$ **do**
  $x_i \leftarrow S_i.regions[0]$
  **foreach** *seed* $S_j \in \{S_{i+1}, \ldots, S_{\min(N, i+b)}\}$ **do**
    $x_j \leftarrow S_j.regions[0]$
    **if** $x_i > x_j \wedge y_i > y_j \wedge x_i + \ell_i > x_j + \ell_j$ **then**
      $d_x \leftarrow x_i - x_j$
      $d_y \leftarrow y_i - y_j$
      $g \leftarrow |d_x - d_y|$
      `// Putative update is the match score minus the gap penalty`
      $u \leftarrow C[i] + \min\{\min\{d_x, d_y\}, \ell_j\} - \lceil 0.01 \cdot L \cdot g + 0.05 \log_2 g \rceil$
      **if** $u > C[j]$ **then**
        $C[j] = u$ `// Update the score if better`
        $B[j] = i$
`// Backtracking to reconstruct the best scoring chains`
$I \leftarrow$ make_heap$\{0, \ldots, N\}$
$i \leftarrow$ pop_heap$(I)$
$s \leftarrow C[i]$
$\mathcal{C} = \{\}$
**while** *true* **do**
  $\mathcal{C}_{\mathrm{cur}} = \{\}$
  **while** $i > 0$ **do**
    $\mathcal{C}_{\mathrm{cur}} \leftarrow \{S_i\} \oplus \mathcal{C}_{\mathrm{cur}}$
    $i \leftarrow B[i]$
  $i \leftarrow$ pop_heap$(I)$
  **if** $(C[i] < s) \vee (|\mathcal{C}| \neq |\mathcal{C}_{cur}|) \vee (\exists j \in \{1, \ldots, |\mathcal{C}|\}\ s.t.\ \mathcal{C}[j] \neq \mathcal{C}_{cur}[j])$ **then**
    **break**
  `/* Only merge this chain to the previous one if it has the same score and matches the same regions in the query */`
  **foreach** $j \in \{1, \ldots, |\mathcal{C}|\}$ **do**
    $\mathcal{C}[j].regions \leftarrow \mathcal{C}[j].regions \oplus \mathcal{C}_{\mathrm{cur}}[j].regions$
**return** $(\mathcal{C}, s)$

### 9 Supplemental Figures

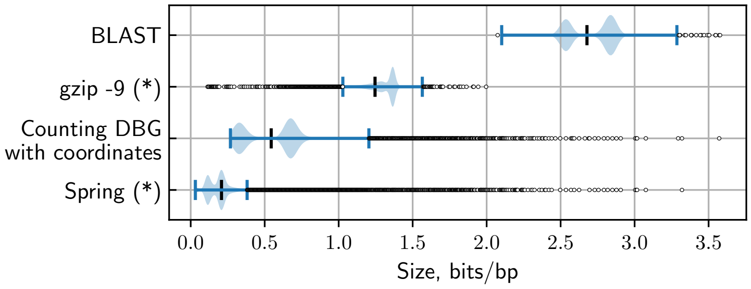


**Supplemental Figure 3.** Compression of Virus PacBio HiFi read sets. – Size distributions for representation of read sets with state-of-the-art indexing and compression methods for all 152,418 virus PacBio HiFi read sets from SRA. BLAST databases were constructed without the m-mer index (MegaBLAST). (*) gzip and Spring do not provide a searchable index and can only be used for compression.

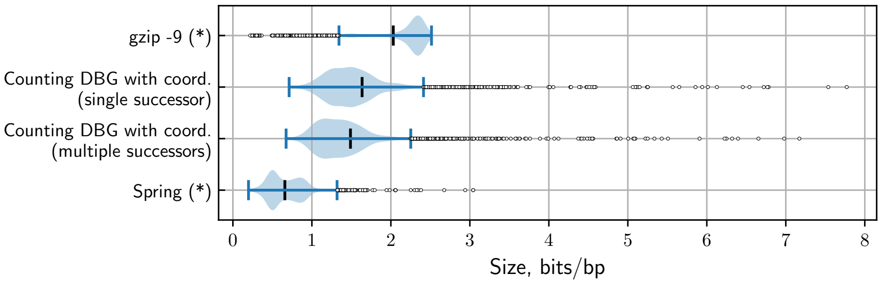


**Supplemental Figure 4.** Compression of Illumina RNA-Seq read sets. – Distribution of compression ratios across the RNA-Seq read sets for five different compression strategies (top to bottom): i) *gzip -9*; ii) our method using only a single successor node; iii) our method using multiple successor nodes; iv) *Spring*. (*) Methods marked with an asterisk do not provide a searchable index and can only be used for compression.

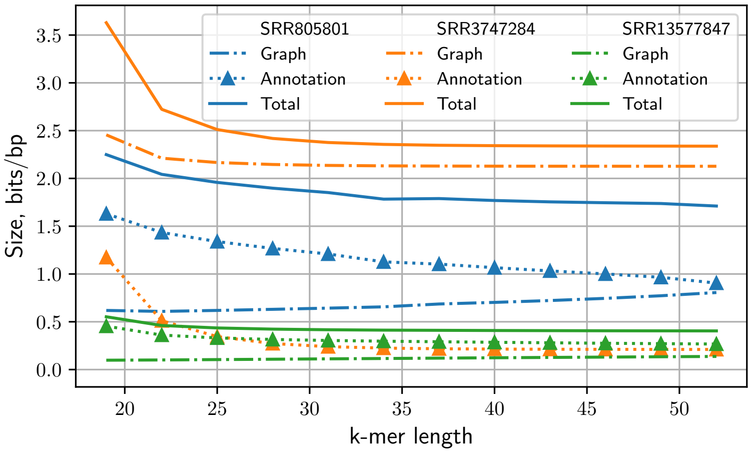


**Supplemental Figure 5.** Index size vs. k-mer length for Illumina (SRR805801), PacBio (SRR3747284), and PacBio HiFi (SRR13577847) reads. – Measurement of the total size of the index (y-axis) for 12 different k-mer length, ranging from 19 to 52 (x-axis). The total index size (solid lines) is broken down into the graph (dash-dot line) and annotation (triangle points) components.
